# Impaired Desynchronization of Beta Activity Underlies Memory Deficits in People with Parkinson’s Disease

**DOI:** 10.1101/667550

**Authors:** Hayley J. MacDonald, John-Stuart Brittain, Bernhard Spitzer, Simon Hanslmayr, Ned Jenkinson

## Abstract

There is a pressing need to better understand the mechanisms underpinning the increasingly recognised non-motor deficits in Parkinson’s disease. Brain activity during Parkinson’s disease is excessively synchronized within the beta range (12–30Hz). However, relatively little is known about how the abnormal beta rhythms impact on non-motor symptoms. In healthy adults, beta desynchronization is necessary for successful episodic memory formation. We investigated whether there was a direct relationship between decreased beta modulation and memory formation in Parkinson’s disease. Electroencephalography recordings were made during an established memory-encoding paradigm. Parkinson’s participants showed impaired memory strength (P = 0.023) and reduced beta desynchronization (P = 0.014) relative to controls. Longer disease duration was correlated with a larger reduction in beta desynchronization, and a concomitant reduction in memory performance. These novel results extend the notion that pathological beta activity is causally implicated in the motor and (lesser appreciated) non-motor deficits inherent to Parkinson’s disease.

## Introduction

Parkinson’s disease (PD) is classified as a movement disorder. However, there is growing recognition that non-motor burdens also significantly impact those suffering with the condition. Non-demented PD patients can experience cognitive difficulties, including long-term memory deficits (for a review see (Raskin, Borod, & Tweedy, 1990; Zgaljardic, Borod, Foldi, & Mattis, 2003) and specifically the ability to recall verbal memory (Cohn, Moscovitch, & Davidson, 2010; Dujardin et al., 2015; Edelstyn et al., 2015).

One striking feature of PD demonstrated repeatedly over the last 20 years is that the electrical activity recorded from basal ganglia (BG) networks in people with PD is excessively synchronized within the beta frequency range (12–30Hz) compared to healthy controls. Under normal circumstances beta activity is modulated with voluntary movement, where the amplitude of oscillations (power) in the beta range drops at the onset of movement and rises again at the end. It is suggested that elevated beta is associated with tonic motor state and event-related desynchronization (ERD) within BG networks “allows” movement to take place (Brittain & Brown, 2014; Joundi et al., 2013), and as such the hyper-synchronized beta state seen in PD prevents desynchronization and thus interferes with voluntary movement (Jenkinson & Brown, 2011). Indeed, therapies that reduce the hyper-synchronized activity, such as dopamine replacement therapy (Ray et al., 2008) or deep brain stimulation (Eusebio et al., 2011), also proportionately improve bradykinesia and rigidity (Ray et al., 2008). Interestingly, beta desynchronization can also occur in the absence of motor output during imagined voluntary movements (McFarland, Miner, Vaughan, & Wolpaw, 2000; Miller et al., 2010). However, to date the link between exaggerated beta activity and motor symptoms in PD remains circumstantial and correlative. It therefore remains an unresolved question as to whether pathological beta activity is causal or an epiphenomenon.

Given that beta activity has shown elevated coupling throughout the BG–thalamocortical circuit in PD, and that this coupling has been observed over broad areas of frontal cortex (Litvak et al., 2011), we postulated that the excessive beta seen in PD should interfere with other neural mechanisms that normally operate within these spatial and temporal domains. Identifying such beta dependent processes and demonstrating a deficit of function in PD would provide further evidence that increased beta is responsible for the motor and non-motor symptoms of the disease. Recent experimental evidence suggests a role for beta oscillations in the encoding of explicit long-term memory. Specifically, a greater amount of beta ERD occurs in the left inferior frontal cortex (IFC) during memory formation of words that are subsequently remembered compared with those that are not (Hanslmayr, Spitzer, & Bauml, 2009; Hanslmayr et al., 2011; Meconi et al., 2016; Meeuwissen, Takashima, Fernandez, & Jensen, 2011; Sederberg, Kahana, Howard, Donner, & Madsen, 2003). This relationship is especially strong if the explicit memory strategy requires semantic processing (Hanslmayr et al., 2009). Memory strategies utilizing semantic processing are examples of deep encoding; when people engage with the meaning of the words e.g. put them into the context of a sentence or make a judgment about whether they relate to living/nonliving entities. Conversely, in shallow encoding an individual only engages with the presented items on a superficial and more perceptual level, as opposed to a cognitive level (Craik & Lockhart, 1972). Examples are detecting whether a presented word contains a specific letter, or whether the first and last letters of the word are in alphabetical order (Otten, Henson, & Rugg, 2001). Unlike in deep encoding, beta ERD during shallow encoding is not predictive of memory performance (Hanslmayr et al., 2009). Furthermore, beta ERD is not seen when similar words are deeply encoded but using non-semantic strategies (Fellner, Bauml, & Hanslmayr, 2013). Therefore, it appears that beta desynchronization is specifically driven by the semantic nature of the encoding task. If the explicit motor deficits in PD are a result of increased beta synchrony in motor areas of the brain, it stands-to-reason that the memory deficits may well be the result of the elevated levels of beta synchrony which prevent the encoding driven ERD required for semantic processing, and memory formation as a result thereof.

Employing a semantic-encoding memory task to investigate the role of pathological beta in PD has several advantages. Firstly, it removes the confound of movement during the beta desynchronization window. Therefore, if a relationship exists between behavior and beta ERD this would argue against impaired beta desynchronization seen in the motor system being an epiphenomenon that merely reflects the paucity of movement in people with PD. Secondly, semantic processing (Gabrieli, Poldrack, & Desmond, 1998) and episodic memory formation (Otten & Rugg, 2001) recruit the *left* IFC. This is important since dynamic modulation of beta has already been shown to be compromised in PD within the cortical-BG network including *right* IFC and subthalamic nucleus (STN) (Brittain et al., 2012; Swann et al., 2011; Swann et al., 2009). Given the coherent beta activity within cortico-BG circuitry (Hirschmann et al., 2011; Litvak et al., 2011) and bidirectional communication (Horschig et al., 2015; Lalo et al., 2008) within these circuits, we would predict that pathological beta would equally affect *left* IFC beta desynchronization and therefore impair episodic memory that recruits semantic encoding strategies. Intriguingly, it has been demonstrated behaviourally that PD patients do show a specific memory deficit when recollecting deep-encoded words, but no deficit in shallow-non-semantic encoding (Cohn et al., 2010). If this specific deficit can be shown to be associated with the inability to sufficiently desynchronize beta activity, it would demonstrate that impaired modulation of beta might underlie at least some of the higher cognitive symptoms associated with the disease. Finally, we have demonstrated a causal relationship between beta power desynchronization in left inferior prefrontal cortex and memory performance in young healthy adults (Hanslmayr, Matuschek, & Fellner, 2014). Elucidating a direct relationship between beta power ERD and episodic memory performance in PD would therefore strongly argue for a causal role of hyper-synchronized beta oscillations in the symptoms of PD.

Given this background, the current study aimed to determine whether there is a direct relationship between impaired beta ERD and the long-term memory deficits observed in non-demented PD. The study design, hypotheses and analyses were pre-registered (MacDonald H, Jenkinson N, Hanslmayr S. Memory encoding and beta desynchronisation in Parkinson’s disease [Internet]. 2016 Available from: https://osf.io/vb64n/). We recorded surface electroencephalography (EEG) during an established memory-encoding paradigm to examine beta oscillations in PD patients and healthy controls during deep-semantic and shallow-non-semantic encoding. We hypothesized that PD patients would exhibit impaired memory performance compared to healthy controls following deep-semantic encoding but that there would be no difference in memory performance between groups following shallow-non-semantic encoding. We further hypothesized that PD patients would show reduced beta ERD during deep-semantic encoding compared to healthy controls, but that there would be no difference in desynchronization between groups during shallow-non-semantic encoding.

## Results

### Participants

Twenty nine adults with PD and 34 healthy control adults with no known neurological impairment were recruited into the study from local PD community groups and research volunteer databases. This pre-registered recruitment target (see https://osf.io/vb64n/) was calculated to account for 10 % drop out and that some participants might be unable to adequately perform the memory task (e.g. insufficient number of remembered items) while still being sufficient to detect a large behavioural effect size (Cohn et al., 2010: Experiment 1) and obtain a power of 0.9. Data for 3 control participants were removed due to not being able to perform the memory task correctly, and for 1 PD participant due to a change in diagnosis. Demographic information for the remaining 31 control and 28 PD participants is provided in Table 1. Patients were at an average disease duration of 6 ± 4 years (range 0.3–14) and tested on their normal medications to avoid the confound of exacerbated motor symptoms. See Table 2 for demographic and clinical data for each individual PD participant. All participants were native English speakers, had completed education at secondary or tertiary level, had no history of dementia, had normal or corrected-to-normal vision and completed the Oxford Cognitive Screen Plus questionnaire (Demeyere et al., 2016) as an assessment of global cognitive function. The two groups did not differ with respect to age, global cognitive function, or level of education (all *P* > 0.254). All results are shown as group means ± standard error.

**Table 1.** Participant demographics and global cognitive function. Values are mean (standard deviation) unless otherwise specified. HC: healthy controls; PD: Parkinson’s disease; OCS-Plus: Oxford Cognitive Screen Plus questionnaire (max 10). Education is grouped into 1: no formal education; 2: primary school; 3: secondary school; 4: tertiary level.

|  | **HC** | **PD** |
| --- | --- | --- |
| **Age (y)** | 67 (9) | 65 (6) |
| **Education** | 3.8 (0.4) | 3.9 (0.4) |
| **Gender** | 13F/18M | 8F/20M |
| **Disease Duration (y)** | N/A | 6 (4) |
| **Handedness** | 3L/28R | 5L/23R |
| **OCS-Plus** | 9.7 (0.5) | 9.7 (0.5) |

**Table 2.**
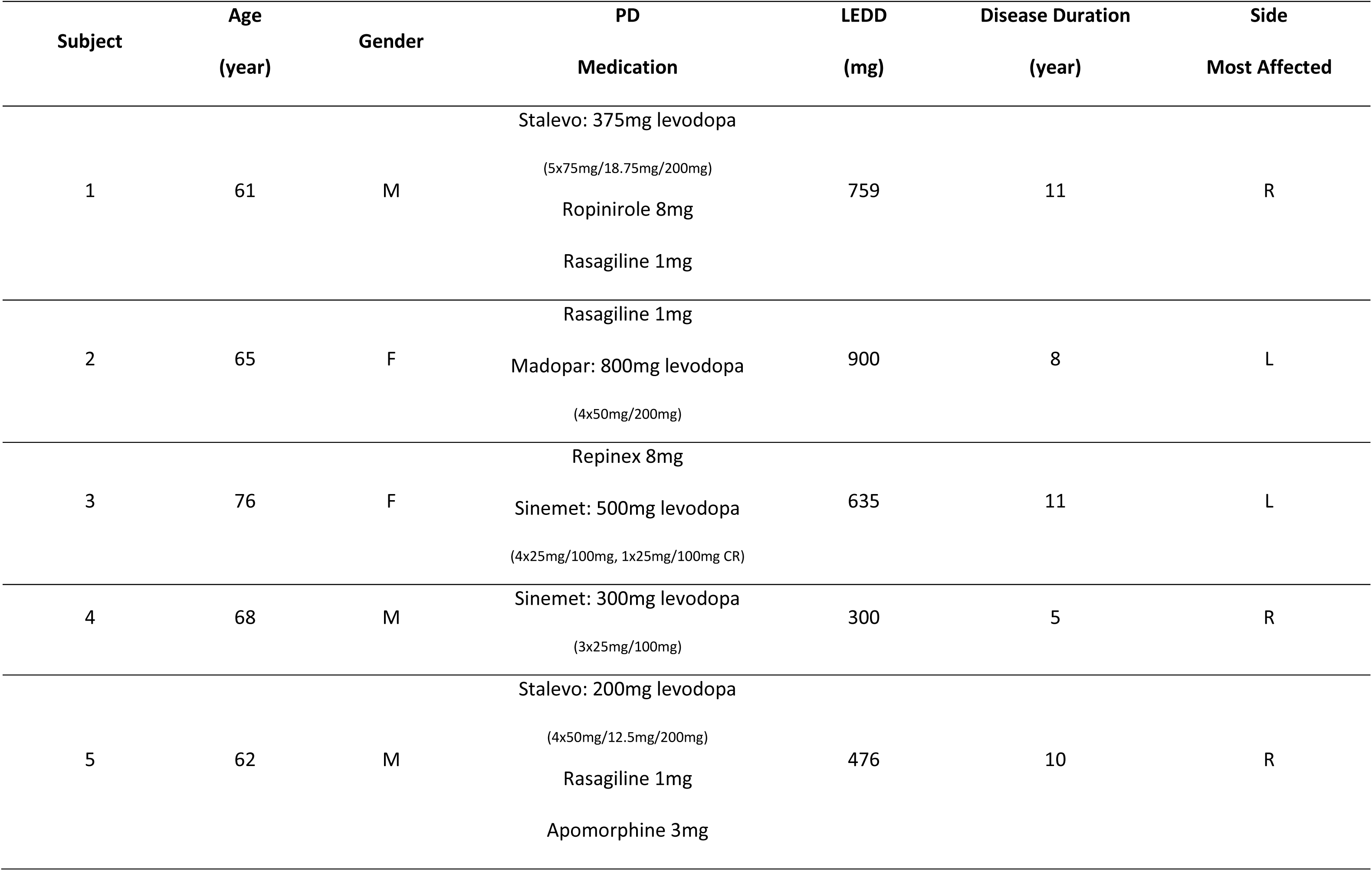
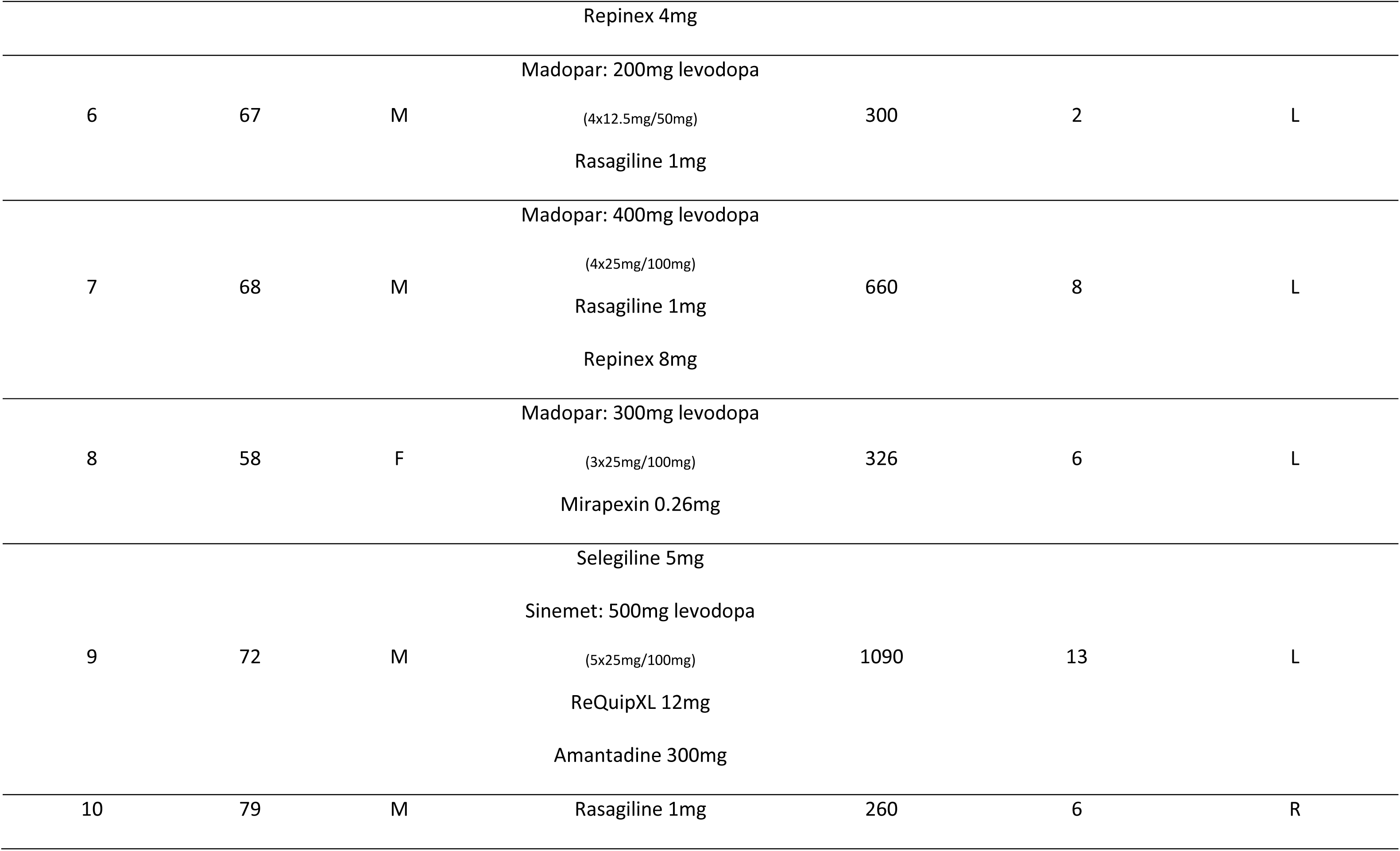
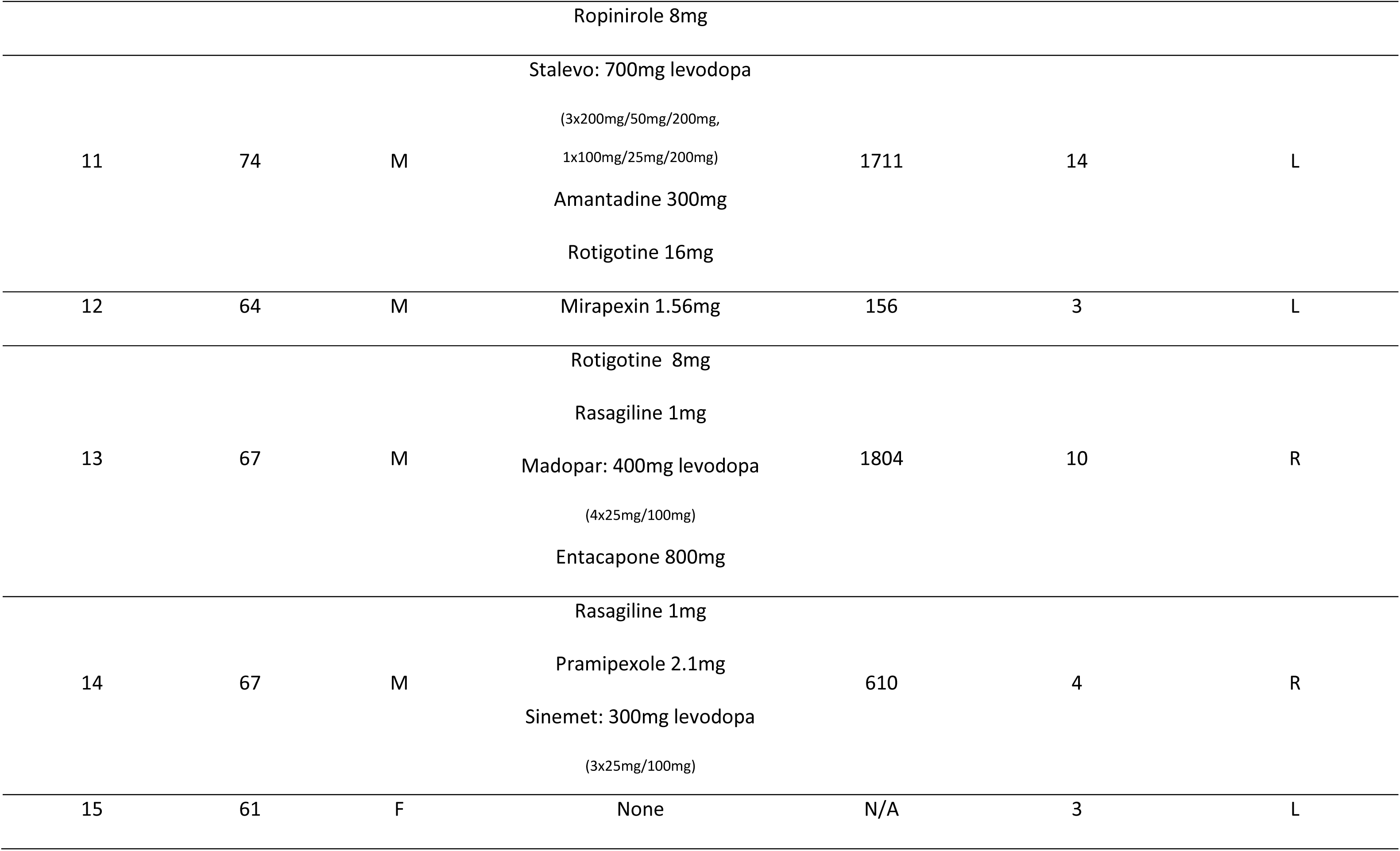
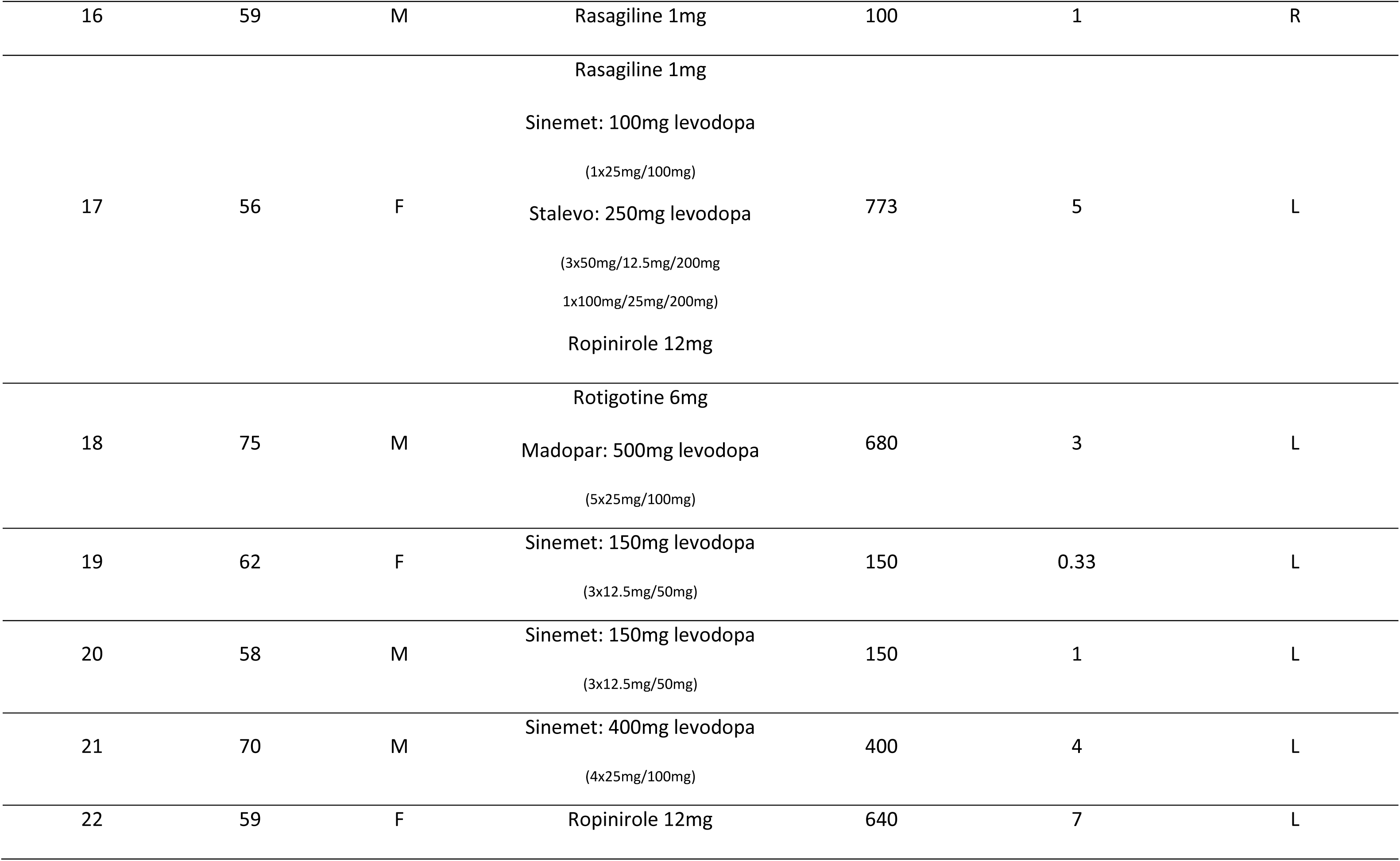
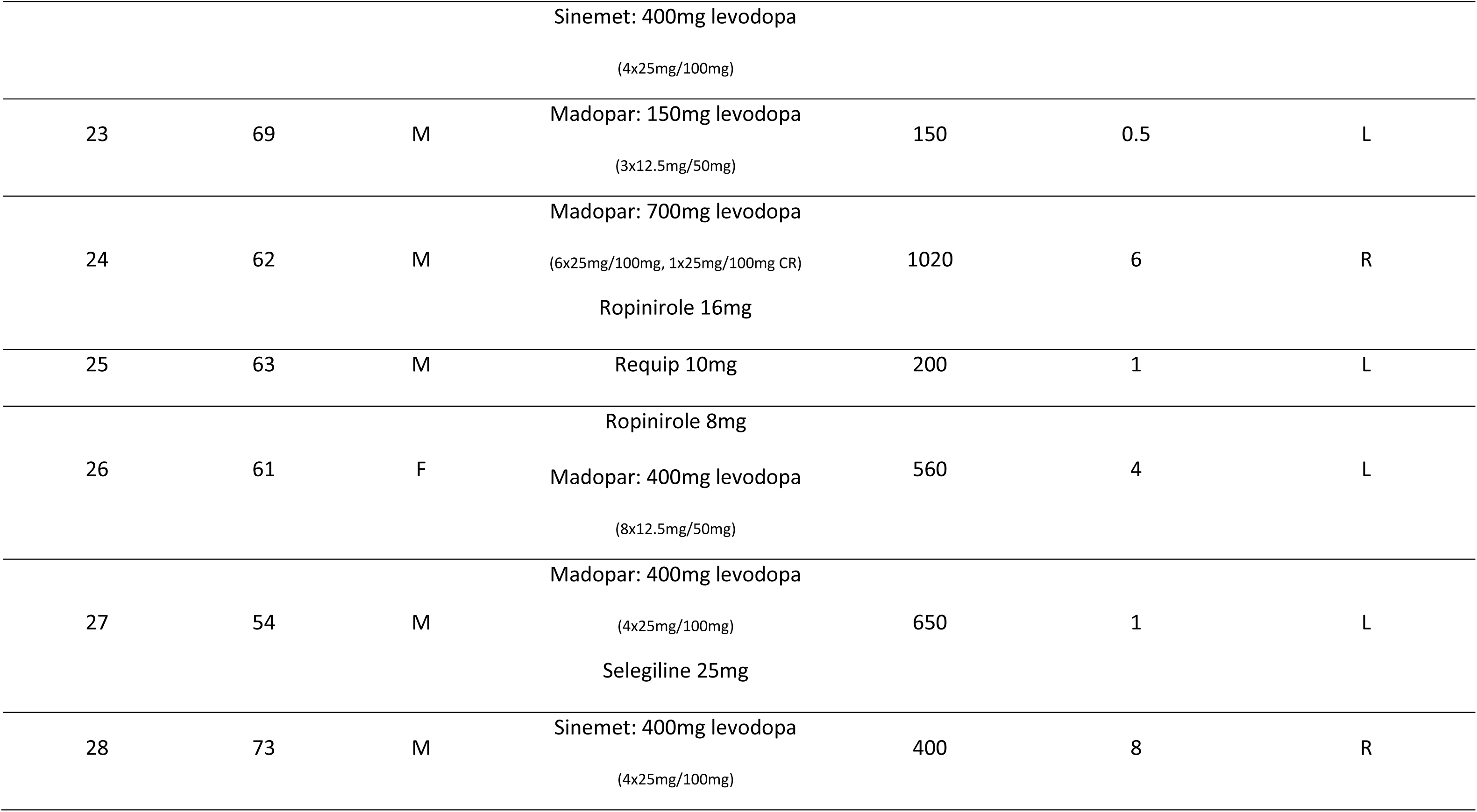
Demographic and clinical data for Parkinson’s participants. PD: Parkinson’s disease; LEDD: levodopa equivalent daily dose; CR: continuous release.

### Behavioural

#### Memory strength

In the deep-semantic encoding blocks, participants judged whether the presented word was animate i.e. whether it referred to the property of a living entity. In the shallow-non-semantic encoding blocks, participants judged whether the first and last letters of the word were in alphabetical order. These encoding instructions have been used previously to investigate subsequent memory effects (Hanslmayr et al., 2009; Otten & Rugg, 2001). Recognition testing at the end of each block required participants to rate their confidence as to whether a word presented was one encountered during encoding, or was a new word. This recognition stage was used to calculate memory strength.

Normal distributions were confirmed for all behavioural data sets (all *P* > 0.423). A mixed-effects repeated measures (RM) ANOVA on memory strength (d’) revealed no main effect of Group (F_1,57_ = 2.494, *P* = 0.120) but a main effect of Encoding (F_1,57_ = 183.499, *P* < 0.001). Memory performance improved in both groups with the semantic processing strategy associated with deep encoding (2.524 ± 0.105) leading to greater memory strength (d’) during recognition testing compared to shallow encoding (1.249 ± 0.057). There was a Group X Encoding interaction (F_1,57_ = 4.885, *P* = 0.031, Fig. 1A). One-tailed post-hoc *t*-tests revealed no difference in memory strength between groups following shallow-non-semantic encoding (t_57_ = 0.130, *P* = 0.500) but deep-semantic encoding lead to greater memory strength in control participants (2.739 ± 0.145) compared to PD (2.309 ± 0.153; t_57_ = 2.042, *P* = 0.023). Although both groups demonstrated memory benefits from the semantic processing required during deep encoding, controls benefited to a greater degree than PD participants.

**Figure 1.**
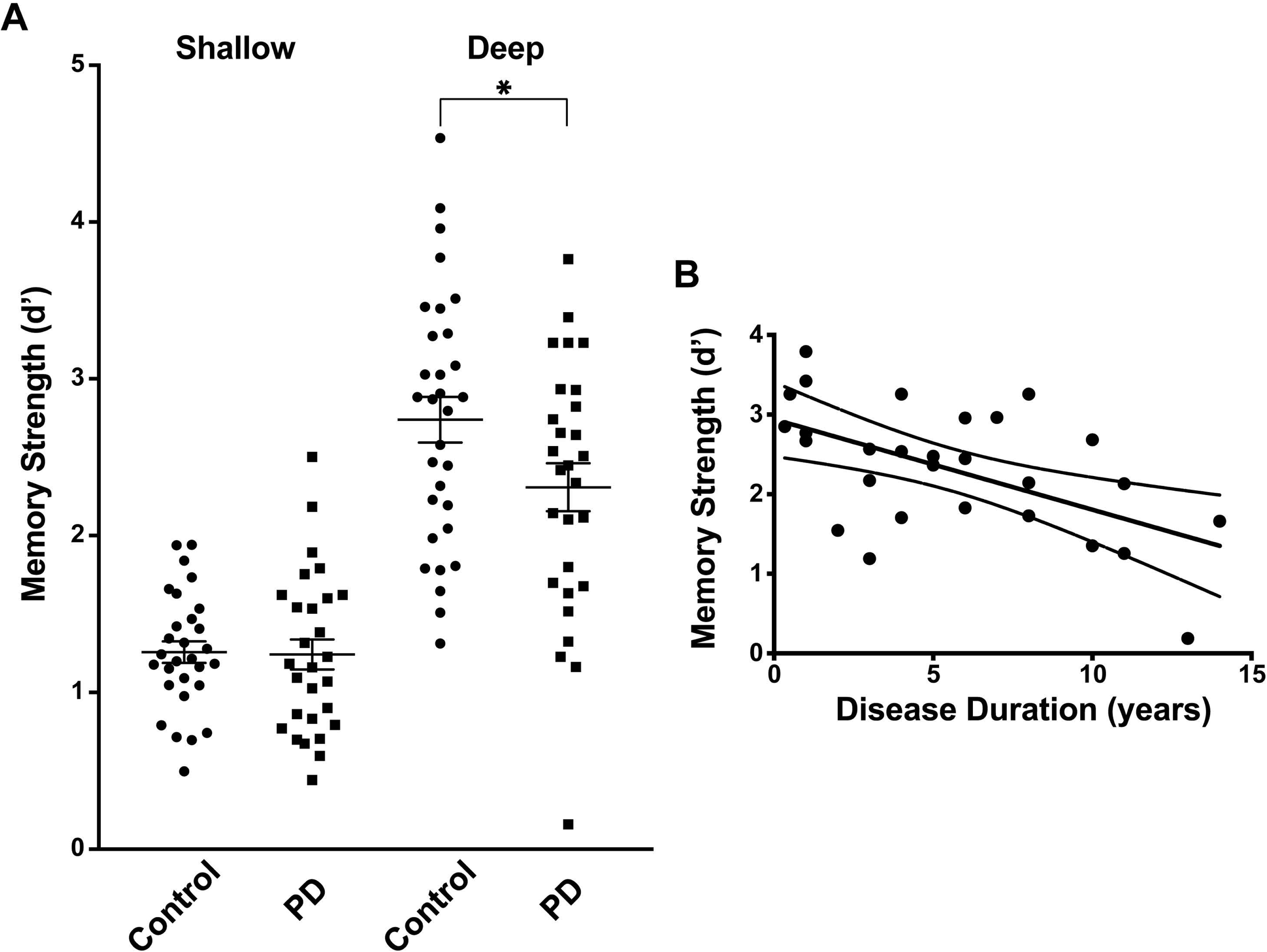
Memory performance. A) Memory performance during encoding conditions illustrating greater memory strength during deep-semantic encoding for healthy controls (N = 31) compared to Parkinson’s disease (PD) participants (N = 28). Error bars denote standard error of the mean. * *P* < 0.05. B) Correlation between deep-semantic encoding memory performance and disease duration for PD participants (*P* = 0.002).

When controlling for age, disease duration had a specific detrimental effect on mechanisms underlying memory formation when semantic processing was required in deep encoding. A LASSO regression was run for PD participants to correlate disease duration with deep-semantic and shallow-non-semantic memory strength as well as age. Only memory strength in the deep-semantic encoding condition was significantly correlated with disease duration (Fig. 1B, F_1,27_ = 11.533, *P* = 0.002, other *P* > 0.242). A similar regression analysis to correlate age and memory strength in controls was not performed as the assumption of normality was violated for age.

#### Encoding reaction time and accuracy

Reaction times and response accuracies were recorded during the *encoding stage* when participants were responding ‘yes’ or ‘no’ with button presses in response to deep-semantic and shallow-non-semantic judgements.

For reaction time, a mixed-effects RM ANOVA produced a main effect of Encoding (F_1,55_ = 6.430, *P* = 0.014) but no effect of Group (F_1,55_ = 1.289, *P* = 0.261) or Encoding X Group interaction (F_1,55_ = 0.764, *P* = 0.386). For both groups, reaction time was faster in shallow-non-semantic encoding (1.12 ± 0.03 s) compared to deep-semantic encoding (1.17 ± 0.03 s) by an average of 50 ms. Similarly, for accuracy, there was a main effect of Encoding (F_1,55_ = 139.156, *P* < 0.001) but no effect of Group (F_1,55_ = 0.044, *P* = 0.834) or Encoding X Group interaction (F_1,55_ = 0.119, *P* = 0.732). Accuracy was higher in shallow-non-semantic encoding (90.9 ± 1.0 %) compared to deep-semantic encoding (75.2 ± 1.1 %) for both groups as expected. The lack of any main effects or interactions with group indicate the significant difference in memory strength between groups in the deep-semantic condition is therefore unlikely to be driven by perceptual differences during encoding.

### EEG

All EEG analysis and presented data are from the *encoding stage*. EEG data from 1 control and 2 PD participants could not be used due to technical problems or large movements from dyskinesia, leaving 30 control and 26 PD EEG data sets for analysis. In alignment with previous EEG studies, and as per our pre-registered protocol, post-stimulus beta power decreases are expected to be associated with successful memory formation in healthy (Hanslmayr et al., 2009; Hanslmayr et al., 2011) and patient populations (Meconi et al., 2016). Therefore lower beta from 12 – 20 Hz was the main frequency range of interest for all dependent measures (see https://osf.io/vb64n/) over 0 – 1.5s relative to stimulus onset (i.e. word presentation).

As hypothesised, the cluster-based permutation testing on all electrodes showed that controls demonstrated greater beta ERD during deep-semantic encoding of subsequently remembered words (Hits) compared to PD participants (cluster stat = −150.1, *P* = 0.014, Fig. 2A & B show beta ERD for electrodes in significant cluster), however no difference between groups emerged during shallow-non-semantic encoding (cluster stat = −3.7, *P* = 0.326, Fig. 2C & D show beta ERD for electrodes in largest cluster that did not reach significance). A mixed-effects RM ANOVA on averaged beta (over 0 – 1.5 s, 12 – 20 Hz) further supported this finding by producing a significant Encoding X Group interaction (F_1,54_ = 6.959, *P* = 0.011) that confirms the difference between groups in deep-semantic encoding (t_54_ = 2.910, *P* = 0.005) is significantly different to shallow-non-semantic encoding (t_54_ = 1.030, *P* = 0.307). There were no main effects of Encoding (F_1,54_ = 0.612, *P* = 0.437) or Group (F_1,54_ = 3.946, *P* = 0.052). Therefore, a difference in beta ERD between groups is seen only in the deep-semantic encoding condition, indicating that there is an ERD deficit in the PD group that occurs specifically during deep-semantic processing.

**Figure 2.**
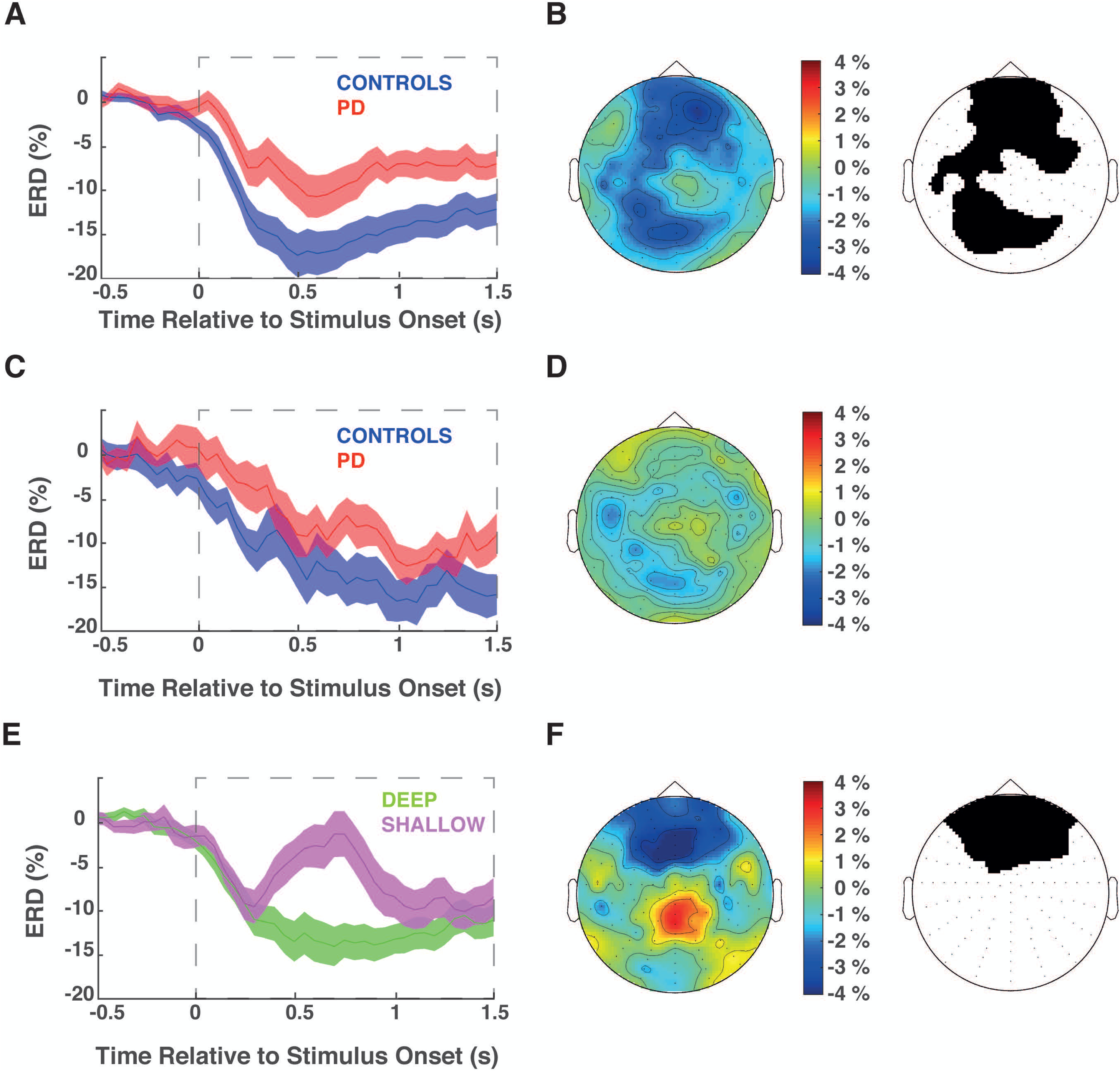
Event related desynchronization. Average beta (12 – 20 Hz) event related desynchronization (ERD) for electrodes in significant and/or largest cluster identified during cluster-based statistical analysis. Top row: between group differences during deep-semantic encoding of remembered words; middle row: between group differences during shallow-non-semantic encoding of remembered words; bottom row: differences within healthy participants between deep-semantic and shallow-non-semantic encoding of remembered words. Grey dashed squares indicate time window used in statistical analysis to identify significant electrode clusters over 12 – 20 Hz. Time course of beta ERD averaged over electrodes contributing to significant and/or largest cluster during encoding of subsequently successfully remembered words for controls (blue, N = 30) compared to Parkinson’s disease (PD) participants (red, N = 26) in the deep-semantic encoding (A) and shallow-non-semantic encoding (C) conditions. A power decrease is denoted with negative values. Only deep-semantic encoding showed a significant difference between groups (electrodes contributing to significant cluster black in panel B). Topographical maps show the location of the ERD differences between groups in deep-semantic (B) and shallow-non-semantic (D) encoding, with colder colours indicating significantly greater ERD in controls compared to PD participants. Cluster shown for shallow-non-semantic encoding in C and D did not reach significance. E) Time course of beta ERD averaged over electrodes contributing to significant cluster during encoding of subsequently successfully remembered words for deep-semantic (green) compared to shallow-non-semantic encoding (magenta) in controls. A power decrease is denoted with negative values. Only controls showed a significant difference between encoding conditions (electrodes contributing to significant cluster black in F). Topographical map in F shows the location of ERD differences between encoding conditions, with colder colours indicating significantly greater ERD in deep-semantic compared to shallow-non-semantic encoding. No cluster identified between encoding conditions for PD patients.

The relationship between beta ERD and the deep-semantic encoding condition is reinforced by the similar pattern of beta ERD seen during the encoding of words that were not successfully remembered (Misses). Misses in controls were associated with greater beta ERD during deep-semantic encoding when compared to PD participants (cluster stat = −54.1, *P* = 0.031), however no difference between groups emerged during shallow-non-semantic encoding (cluster stat = −3.8, *P* = 0.330). A mixed effects RM ANOVA similarly produced main effects of Encoding (F_1,54_ = 5.450, *P* = 0.023) and Group (F_1,54_ = 6.155, *P* = 0.016) and a significant Encoding X Group interaction (F_1,54_ = 5.975, *P* = 0.018). The interaction confirms the difference between groups in deep-semantic encoding (t_54_ = 3.367, *P* = 0.001) is significantly different to shallow-non-semantic encoding (t_54_ = 0.919, *P* = 0.362). The fact that a difference in beta ERD is seen between groups during encoding of both remembered and forgotten items implies the difference is related to deep-semantic encoding in general. This overall reduced beta desynchronization may lead to reduced memory performance in PD participants.

Successful memory formation specifically involving deep-semantic processing was associated with greater beta ERD. Within groups, controls demonstrated greater beta ERD for subsequently remembered words during deep-semantic compared to shallow-non-semantic encoding (cluster stat = −94.4, *P* = 0.012, Fig. 2E & F show beta ERD for electrodes in significant cluster). Interestingly at a group level, PD participants did not show significantly greater ERD in deep-semantic encoding compared to shallow-non-semantic (no significant clusters were identified), although they did show a behavioural benefit of deep-semantic encoding, albeit to a lesser extent than controls. Based on findings of left IFC beta being specifically linked to memory strength in healthy controls (Hanslmayr et al., 2009; Hanslmayr et al., 2011; Meeuwissen et al., 2011), we did an additional correlational analysis focusing on left frontal beta in PD patients. Despite no group-level effect, linear regressions illustrated that PD participants who showed greater beta ERD over left frontal electrodes also had significantly greater memory strength during deep-semantic encoding (*P* = 0.008, R^2^ = 0.256, Fig. 3A) but that disease duration negatively correlated with left frontal maximum beta ERD (P = 0.007, R^2^ = 0.263, Fig. 3B). PD participants earlier in the disease who were able to achieve greater beta ERD in left frontal electrodes benefited more from deep-semantic encoding strategies of memory formation.

**Figure 3.**
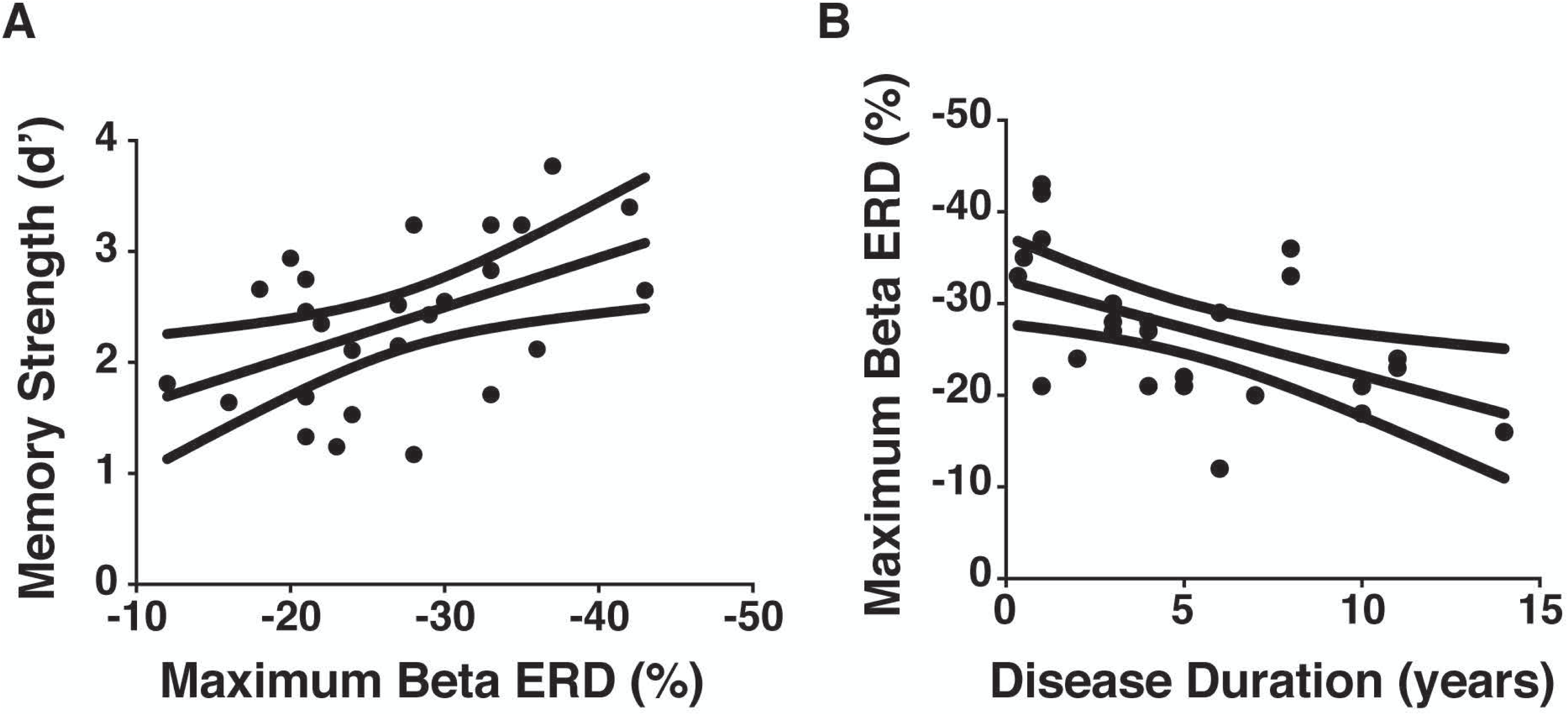
Correlations in Parkinson’s disease patients. A) Correlation between deep-semantic encoding memory performance and maximum beta ERD over left frontal electrodes for PD participants (N = 26, *P* = 0.008, R^2^ = 0.256). B) Correlation between maximum beta ERD over left frontal electrodes and disease duration for PD participants (N = 26, P = 0.007).

The secondary dependent measure was the subsequent-memory effect (SME) in beta power which compared power between high confidence hit (i.e. subsequently strongly remembered) and miss (i.e. subsequently forgotten) trials (Brewer, Zhao, Desmond, Glover, & Gabrieli, 1998; Hanslmayr et al., 2009; Otten et al., 2001). This categorization in the *encoding stage* depended on the participant’s response in the *recognition stage* and their individualized receiver operating characteristic (ROC) curves (Hanslmayr *et al*. 2009; see Materials and Methods). The SME results broadly replicated a number of previous findings (Hanslmayr et al., 2009; Hanslmayr et al., 2011; Meconi et al., 2016) and further support the importance of beta ERD as the mechanism underlying successful memory formation through deep-semantic encoding strategies: there was a significant SME in deep-semantic encoding for controls (cluster stat = −42.2, *P* = 0.027, Fig. 4A & B illustrate beta ERD for electrodes in significant cluster) and a SME approaching significance for PD participants (cluster stat = −22.3, *P* = 0.097, Fig. 4C & D illustrate beta ERD for electrodes in the largest cluster).

**Figure 4.**
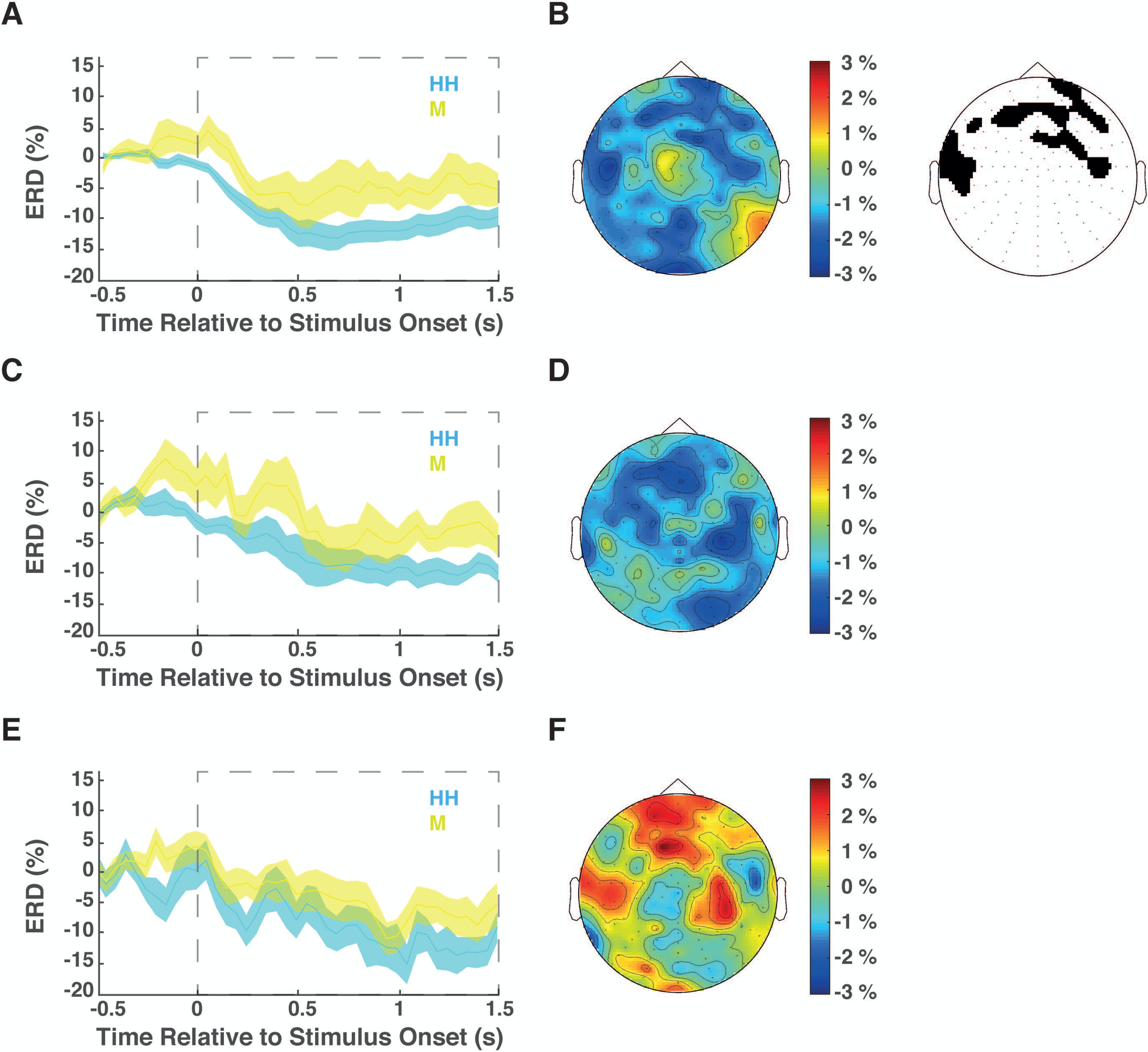
Subsequent memory effects. Average beta (12 – 20 Hz) event related desynchronization (ERD) for electrodes in significant and/or largest cluster identified during cluster-based statistical analysis. Top row: differences within healthy participants between remembered and forgotten words during deep-semantic encoding; middle row: differences within PD patients between remembered and forgotten words during deep-semantic encoding; bottom row: differences within healthy participants between remembered and forgotten words during shallow-non-semantic encoding. Grey dashed squares indicate time window used in statistical analysis to identify significant electrode clusters over 12 – 20 Hz. Time course of beta ERD averaged over electrodes contributing to significant and/or largest cluster during high confidence hit (HH, cyan) compared to miss (M, yellow) trials in deep-semantic encoding for controls (A, N = 30) and PD participants (C, N = 26). Both groups demonstrated greater ERD during encoding of subsequently remembered (HH) compared to forgotten (M) words, but only the cluster in controls reached significance (electrodes contributing to significant cluster black in B). Topographical maps show the location of the ERD differences between words in deep-semantic encoding for controls (B) and PD patients (D), with colder colours indicating greater ERD for remembered compared to forgotten words. Time course (E) and location (F) of beta ERD averaged over electrodes contributing to largest, non-significant cluster during high confidence hit (HH, cyan) compared to miss (M, yellow) trials in shallow-non-semantic encoding for controls (N = 30). No cluster identified between remembered and forgotten words in shallow-non-semantic encoding for PD patients.

Importantly, there was no significant SME associated with shallow-non-semantic encoding (controls: cluster stat = −2.1, *P* = 0.698, Fig. 4E & F illustrate beta ERD for electrodes in largest cluster that did not reach significance; PD: no clusters were identified). A mixed-effects RM ANOVA showed a main effect of Encoding (F_1,54_ = 24.265, *P* < 0.000), confirming that deep-semantic encoding produced a greater average SME (−6 ± 1 %) compared to shallow-non-semantic encoding (1 ± 0.7 %). There was no main effect of Group (F_1,54_ = 0.007, *P* = 0.935) or Encoding X Group interaction (F_1,54_ = 0.023, *P* = 0.880). The lack of an interaction was expected as, although PD participants remembered fewer items than controls following deep-semantic encoding, the remembered items in both groups should be accompanied by similar electrophysiological signatures (i.e. SME) as in both cases they lead to the same behavioural outcome – that of remembering (i.e. d’ above zero).

## Discussion

The study confirmed our pre-registered hypotheses and produced several novel findings that provide the first evidence of impaired beta modulation being associated with a non-motor symptom of PD. PD participants showed impaired memory strength compared to healthy controls but only following deep-semantic encoding of words. This behavioural finding was mirrored by the EEG results which demonstrated that PD participants exhibited reduced beta ERD compared to healthy controls but again only during deep-semantic memory formation. Furthermore, a correlation between disease duration and an increased deficit in deep-semantic encoding suggested the neuropathology of PD has a specific detrimental effect on the mechanisms underlying deep-semantic memory formation leading to both reduced beta ERD and reduced memory strength. This is reinforced by that fact that participants with PD who showed greater beta ERD over left frontal electrodes benefited to a greater extent from the deep-semantic encoding memory strategy. There were no differences between the groups in age, global cognitive function, education or perception during encoding that could explain these behavioural or EEG results. Therefore, our results appear to be specific to episodic memory formation as a result of deep-semantic processing. Overall, our findings strengthen the idea that dysfunctional beta oscillations are likely to be the cause of PD symptoms in both motor and non-motor domains.

Parkinson’s disease did not cause impaired memory performance in general, but rather a specific deficit in deep-semantic encoding of memory. Deep-semantic encoding in the context of the current study utilized general knowledge about the word to form an abstract representation and evaluate the representation as animate or inanimate. Age-related memory decline is a widely acknowledged fact that is seen across several subdomains, including episodic memory (e.g. see (Shing et al., 2010)). Over and above the aging-related decline, a further decline in episodic memory resulting from deep-semantic encoding appeared to be caused by the mechanisms underlying PD. Replicating previous findings, PD participants were able to employ the non-semantic encoding strategy to build a memory trace of equivalent strength to controls (Cohn et al., 2010). The difference in memory performance between groups was only elucidated following a deep-semantic encoding instruction. In contrast to Cohn and colleagues (Cohn et al., 2010), the current PD participants still showed a behavioural benefit from the deep-semantic encoding memory strategy and those who were less progressed in the disease benefited to a greater degree. People with PD struggle to spontaneously implement the optimal memory encoding strategy (Knoke, Taylor, & Saint-Cyr, 1998). However with explicit encoding instructions, PD participants managed to improve memory with the optimal deep-semantic encoding strategy, albeit to a lesser degree than controls. This finding suggests they are able to recruit the neural mechanisms to process semantic information about the words in the deep encoding condition, but something prevents the formation of a robust memory trace. Overall, people with PD exhibited a limited deep-semantic processing capacity during memory encoding rather than a general deficit in recognition memory.

The deficit in episodic memory performance following a deep-semantic encoding strategy displayed by PD participants was associated with a reduced dynamic range of beta ERD during encoding. Brain oscillations are considered one of the core neural mechanisms for storage and retrieval of long-term memories (Buzsaki & Draguhn, 2004; Fell & Axmacher, 2011) and the extent of neural desynchronization is thought to relate to the degree of information stored in the brain (Hanslmayr, Staudigl, & Fellner, 2012). In the current study, the greater level of beta desynchronization for deep-semantic versus shallow-non-semantic encoding, and words that were subsequently remembered compared to those that weren’t, further supports the importance of beta ERD as the mechanism underlying successful deep-semantic memory formation (Hanslmayr et al., 2009; Hanslmayr et al., 2011; Meconi et al., 2016; Meeuwissen et al., 2011; Sederberg et al., 2007). As both groups displayed similar behavioural outcomes of deep-semantic encoding (i.e. d’ values above zero, although PD participants remembered fewer items than controls), it is not surprising that both groups displayed similar electrophysiological differences between high confidence hits and misses (i.e. a SME). Importantly however, overall beta ERD was significantly reduced in PD participants compared to controls during deep-semantic processing, but not for words encoded with a shallow-non-semantic strategy. This distinction implies that a reduced capacity to decrease beta power following stimulus presentation for PD participants reduced the richness of semantic information encoded in the brain and therefore weakened the memory strength, leading to fewer successfully recognized words and a lower d’ value.

It has been proposed that the relative change in pre-to post-stimulus power is most important for memory performance, rather than absolute power levels (Klimesch, Doppelmayr, & Hanslmayr, 2006; Klimesch, Sauseng, & Gerloff, 2003). PD participants demonstrated decreases in the reactivity of their event-related beta power and therefore reduced encoding capacity. PD participants who were further progressed in the disease demonstrated further reductions in both beta reactivity and memory strength. A reduced dynamic range of BG-thalamocortical beta power in PD can therefore interfere with other neural mechanisms that operate in the beta frequency range apart from movement, including memory formation.

The neural changes causing episodic memory deficits in PD may be the same as those underlying motor symptoms. Memory formation recruits an extensive network of mainly left-lateralized regions for verbal material. This network includes the anterior temporal lobe for storage of conceptual representations and processing concepts at an abstract level (Jefferies & Lambon Ralph, 2006; Patterson, Nestor, & Rogers, 2007), and the IFC and temporoparietal region for strategic search and control processes that are necessary for semantic processing (Binder, Desai, Graves, & Conant, 2009; Jefferies, 2013; Jefferies & Lambon Ralph, 2006). The extent of beta ERD in left prefrontal cortex (PFC), specifically IFC, has been linked to memory performance (Hanslmayr et al., 2009; Hanslmayr et al., 2011). Function of the PFC is heavily influenced by the integrity of dopaminergic input onto frontostriatal connections.

Therefore, it is not surprising that dopaminergic dysfunction seen in PD leads to impaired IFC function, observed in motor tasks that recruit the right IFC as part of the response inhibition network (Bokura, Yamaguchi, & Kobayashi, 2005; Gauggel, Rieger, & Feghoff, 2004; Obeso et al., 2011; Swann et al., 2011). We have extended these findings to also show impairment during a memory task that has been shown to recruit the left IFC during deep-semantic encoding. Previous studies have highlighted the ability of BG oscillatory activity to influence cortical neuronal oscillations recorded with surface EEG (Chung et al., 2018; Horschig et al., 2015). We therefore propose that the same pathological BG beta mechanism causing the motor symptoms in PD is contributing to the deficit in deep-semantic encoding of memory seen in the current study. This would imply a common neural mechanism may underlie a variety of deficits in PD that involve cortico-BG processes which operate predominantly in the beta frequency range.

It is a matter of speculation as to the cause of altered memory-related beta oscillations within PD. However, there are potential candidate mechanisms that could be contributing to pathological beta within the memory domain. For example, long-term potentiation (LTP) in the hippocampus is proposed as the mechanism of synaptic plasticity playing a key role in the formation of long-term memories (Bliss & Collingridge, 1993). Neural oscillations are thought to shape synaptic plasticity by providing temporal windows for neural firing (Hanslmayr, Staresina, & Bowman, 2016), so the differences in cortical beta oscillations in the current study between PD patients and healthy participants might be linked to LTP-like mechanisms. However, the direct relationship between any one form of synaptic plasticity and a specific frequency range of oscillations or a particular type of memory is still unclear and highly speculative, especially in humans and cortical regions. Intriguingly, LTP-like mechanisms that are altered in the motor areas in people with PD (Kishore, Joseph, Velayudhan, Popa, & Meunier, 2012; Lago-Rodriguez et al., 2016; Suppa et al., 2011) are also suggested to be the mechanism behind the reduced modulation (in PD) of movement-related beta that is normally seen in the sensorimotor area during repetitive practice of arm movements in healthy controls (Moisello et al., 2015; Nelson et al., 2017). Therefore, future studies could investigate whether impaired LTP-like mechanisms are linked to the reduced memory performance and reduced beta modulation seen in PD patients in the current paradigm.

Identifying a common neural mechanism behind the motor and non-motor symptoms of PD has implications for treatment and disease monitoring. There are currently no standard treatment options for mild memory and cognitive problems in PD (i.e. mild cognitive impairment). Applying interventions previously shown to decrease hyper-synchronized beta activity such as deep brain stimulation or dopamine replacement therapy (Eusebio et al., 2011; Ray et al., 2008) should in theory also help with memory deficits caused by the same pathology. Considering the inverse relationship demonstrated in the current study between disease progression and both memory performance and beta ERD, it is feasible that this memory paradigm could be developed as a useful surrogate to measure functional beta reactivity. As such, the paradigm could be used as a new and convenient behavioural test to monitor disease progression, with specific applications in telemedicine.

It is important to note that while we present findings that the neural changes causing episodic memory deficits in PD may resemble those underlying motor symptoms, we do not posit that reduced beta de-synchronisation is the sole deficit that emerges in PD. Nor, in-fact, that there is a single source of beta that homogenises symptomology across domains (Spitzer & Haegens, 2017). Instead, we extend the impact of a deficit that has been identified in the motor domain to other (cognitive) areas. This will likely explain some symptoms well, but not all, and should be a consideration when titrating medications to alleviate different aspects of motor and/or cognitive performance. It is important to make this distinction as we are not claiming that beta observed in the motor system directly influences memory encoding – but that beta in memory-relevant areas is also deficient and, while these rhythms are likely to serve a similar functional role, deficits may indeed be graded across functional areas. Hence, motor deficits and memory deficits may be differentially influenced depending on the underlying pathophysiological state.

There are a few limitations to the current study that should be considered. Firstly, the relationship between beta ERD and the behavioural deficit in the PD group is correlational. However, it is the more parsimonious explanation that a common underlying neurological deficit (i.e. impaired beta desynchronization) causes both motor and memory problems than two unrelated behavioural symptoms producing the same epiphenomenon in the beta system. Furthermore, evidence exists for a causal relationship between the strength of beta desynchronization in left PFC and memory performance (Hanslmayr et al., 2014) so the direct relationship shown in the current study would support a causal role of pathological beta in PD symptomology. Extending the findings from Hanslmayr and colleagues, future studies could use transcranial magnetic stimulation to modulate left prefrontal beta in people with PD and look for a causal influence on their episodic memory performance. Secondly, beta desynchronization also plays a role in memory retrieval (Dujardin, Bourriez, & Guieu, 1994; Duzel et al., 2003) and people with PD are thought to use inefficient retrieval strategies (see (Zakzanis & Freedman, 1999). However using recognition, which is one of the simplest ways to test episodic memory, greatly reduced retrieval demands in our task, e.g. compared to free or cued recall. A retrieval based explanation for our behavioural findings is therefore rather unlikely. Nevertheless, we cannot completely discount the contribution of impaired beta desynchronization during retrieval to the reduced recognition memory performance in our study. Our prior hypothesis and pre-registered protocol focused initially on memory encoding because encoding primarily recruits the IFC, while retrieval recruits parietal regions (Burgess & Gruzelier, 2000; Spitzer, Hanslmayr, Opitz, Mecklinger, & Bauml, 2009; Zion-Golumbic, Kutas, & Bentin, 2010). Due to the dopaminergic modulation of frontostriatal connections discussed previously, we expected pathological BG beta in PD would preferentially affect prefrontal cortical regions.

Despite displaying topographical maps in an effort to show the location of ERD differences between groups, the methods used in the current study cannot be used to form a robust conclusion about spatial differences in beta ERD. The location of beta ERD differences in deep-semantic encoding between patients and healthy participants seemed to indicate a widespread cortical deficit in beta desynchronization in PD patients, which included the left frontal region. This widespread difference is in contrast to, for example, more focal differences in beta ERD for healthy participants between deep-semantic and shallow-non-semantic encoding. However scalp-level EEG has limited spatial resolution. Subsequent studies using magnetoencephalography with a much higher spatial resolution would be needed to investigate these results further. Finally, when considering the generalizability of our results, it is worth noting that the PD patients in the current study were mild to moderately impaired in terms of disease severity. Our study therefore cannot directly speak to the relationship between memory impairments and beta oscillations in severely affected PD patients. However, our findings of an inverse relationship between disease duration and both memory performance and beta desynchronization speaks to a general characterisation that will likely extend (alongside other age-related factors) to those severely impaired patients.

### Conclusion

This study provides the first evidence of impaired beta modulation being associated with a non-motor symptom of PD. PD participants showed impaired memory strength and beta ERD compared to healthy controls during deep-semantic encoding. The neuropathology of PD seemed to have a specific detrimental effect on the mechanisms underlying episodic memory formation in a deep-semantic encoding task leading to both reduced memory strength and reduced beta ERD. We propose that the neural changes causing memory deficits in PD may be the same as those underlying motor symptoms i.e. impaired modulation of beta activity within BG– thalamocortical circuitry. Importantly the decrease in beta modulation shown in our study cannot be explained away as an epiphenomenon that scales with decreased movement in PD. Our findings strengthen the idea that dysfunctional beta oscillations are causal in PD symptomology, and extend their implications to non-motor symptoms of the disease.

## Materials and Methods

The study was approved by the University of Birmingham Research Ethics Committee (ERN_09-528AP20) and written informed consent was obtained from each participant. Data collection was carried out during a single laboratory session for each participant at the University of Birmingham.

### Behavioural task

Participants were seated approximately 1 m from a 19 inch computer monitor. Stimuli were presented in black text against a grey background using the Psychophysics Toolbox extension of Matlab (Brainard, 1997). The task was divided into eight blocks and each block into three stages (Fig. 5).

**Figure 5.**
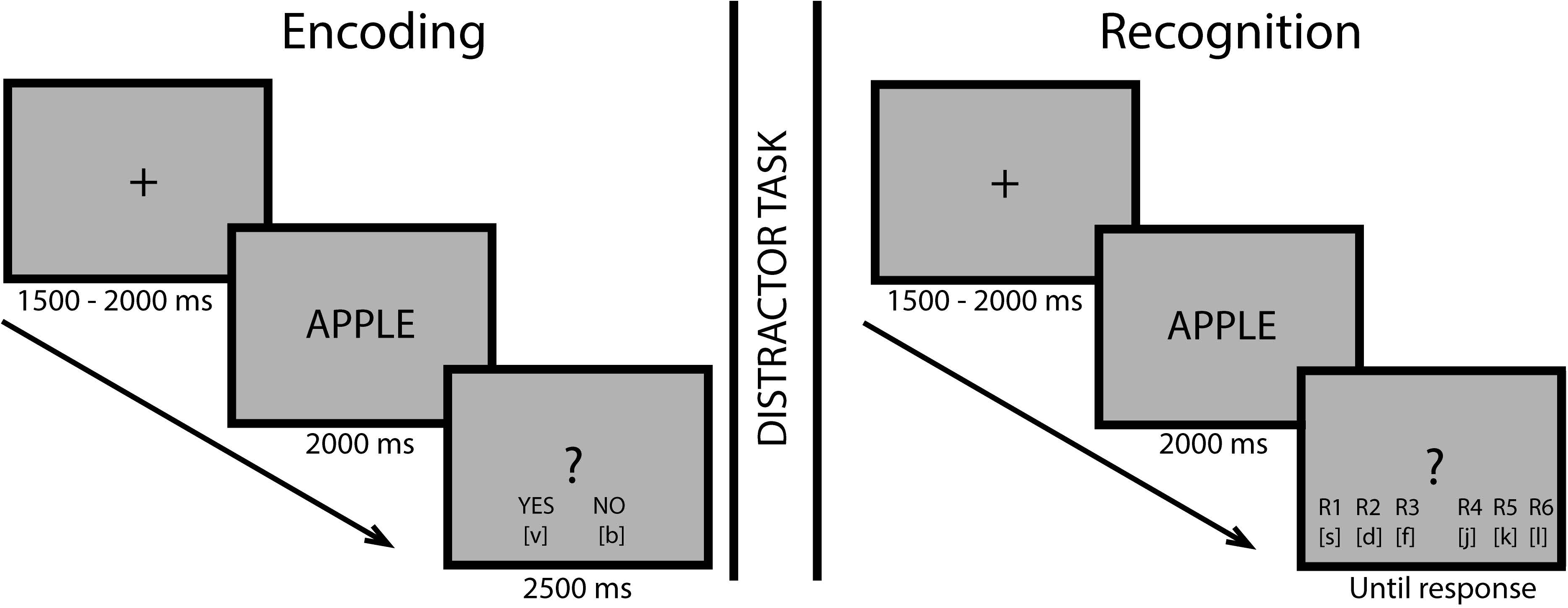
Three stages of memory task. The letters in brackets indicated to participants which button on the keyboard corresponded to which response. In the final screen for a recognition trial participants saw assigned responses (i.e. recollection, very familiar etc.) rather than R1 – R6 which are shown here due to space constraints.

First, there was an *encoding stage*, which required either deep-semantic or shallow-non-semantic encoding of 30 words presented on the screen one at a time. All participants completed four blocks of each encoding. The order of presentation of each encoding-type was counterbalanced across participants. In the deep-semantic encoding blocks, participants judged whether the presented word was animate i.e. whether it referred to the property of a living entity. In the shallow-non-semantic encoding blocks, participants judged whether the first and last letters of the word were in alphabetical order. These encoding instructions have been used previously to investigate subsequent memory effects (Hanslmayr et al., 2009; Otten & Rugg, 2001). Participants responded on each trial by pressing one of two response buttons (“yes” or “no”) on the keyboard using their index and middle finger. PD patients used fingers on their less affected hand and hand assignment was randomized (regardless of hand dominance) across healthy participants for comparison with patients. Button assignment was counterbalanced across patients and participants.

The encoding stimuli were taken from a pool of 240 English words, with a list of 120 per encoding condition selected from the MRC psycholinguistic database (Coltheart, 1981). Encoding lists were matched according to word frequency (10 - 93 per million), concreteness (252 - 593), imageability (452 – 615), number of syllables (1 – 4) and number of letters (3 – 10). Words were randomly drawn from the first encoding list for the first four blocks, and the second list for the last four blocks. The order of encoding instructions rather than encoding lists was counterbalanced across participants. A single trial began with a fixation cross for a variable duration of between 1500 and 2000 ms, followed by word presentation for 2000 ms and ended with a question mark to prompt the participant to respond (for which they were given 2500 ms). Participants were instructed not to react during word presentation but give their response during presentation of the question mark.

The second stage in each block consisted of a distracter task during which 20 faces of famous and non-famous people were presented to the participant one at a time. The participant was required to rate the attractiveness of each face using a 6-point rating scale. The distracter stage was intended to prevent the participants rehearsing the word lists, and also to familiarize participants with the 6-button ratings which were to be used in the subsequent recognition stage.

In the final *recognition stage* of each block, the 30 previously encoded words and 15 novel stimuli words drawn from the same pool were presented to participants one at a time. The order of words was randomized and participants were required to rate their confidence as to whether the word was one encountered in the *encoding stage*, or was a new word. Ratings were given using the 6-point rating scale where response options were R1: recollect, R2: very familiar, R3: familiar, R4: unsure new, R5 sure new, R6: very sure new, using buttons pressed with the index, middle and ring fingers on both hands. The assignment of the buttons was counterbalanced across participants (i.e. R1 – R6 vs R6 – R1), and participants were explicitly instructed to use the full range of confidence ratings. The list of new words was matched to encoding lists for word frequency, concreteness, imageability, number of syllables and number of letters. A trial progressed in the same order and with the same timings as during the *encoding stage*, except that the question mark and button prompts remained on-screen until the participant responded.

### EEG recording

Continuous EEG data were recorded using a 128 channel BioSemi ActiveTwo system (BioSemi) with electrodes positioned at the 128 standard equidistant BioSemi sites. Data were digitized using the BioSemi ActiView software, with a sampling rate of 1024 Hz and filtered between 0.1 and 100 Hz.

### Behavioural data analysis

Reaction times and response accuracies were recorded during the *encoding stage*. Response times were calculated from the onset of the question mark which prompted the participant to respond until button press. Accuracy was calculated as the number of correct Yes or No responses during each type of encoding expressed as a percentage of all words presented for that encoding condition. All other behavioural analysis and presented data are from the *recognition stage*. Trials in the recognition stage were grouped into high confidence hit (HH), low confidence hit (LH) and miss (M) categories, depending on the participant’s response and their individualized receiver operating characteristic (ROC) curves (Hanslmayr *et al*. 2009). Using ROCs enabled objective quantification of individual response biases and corrected for participants’ tendencies to use single buttons of the rating scale differently (Fig. 6). The primary dependent variable, memory strength (d’), was calculated from recognition responses using the following equation.

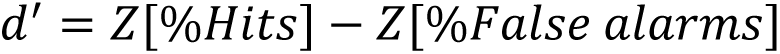

**Figure 6.**
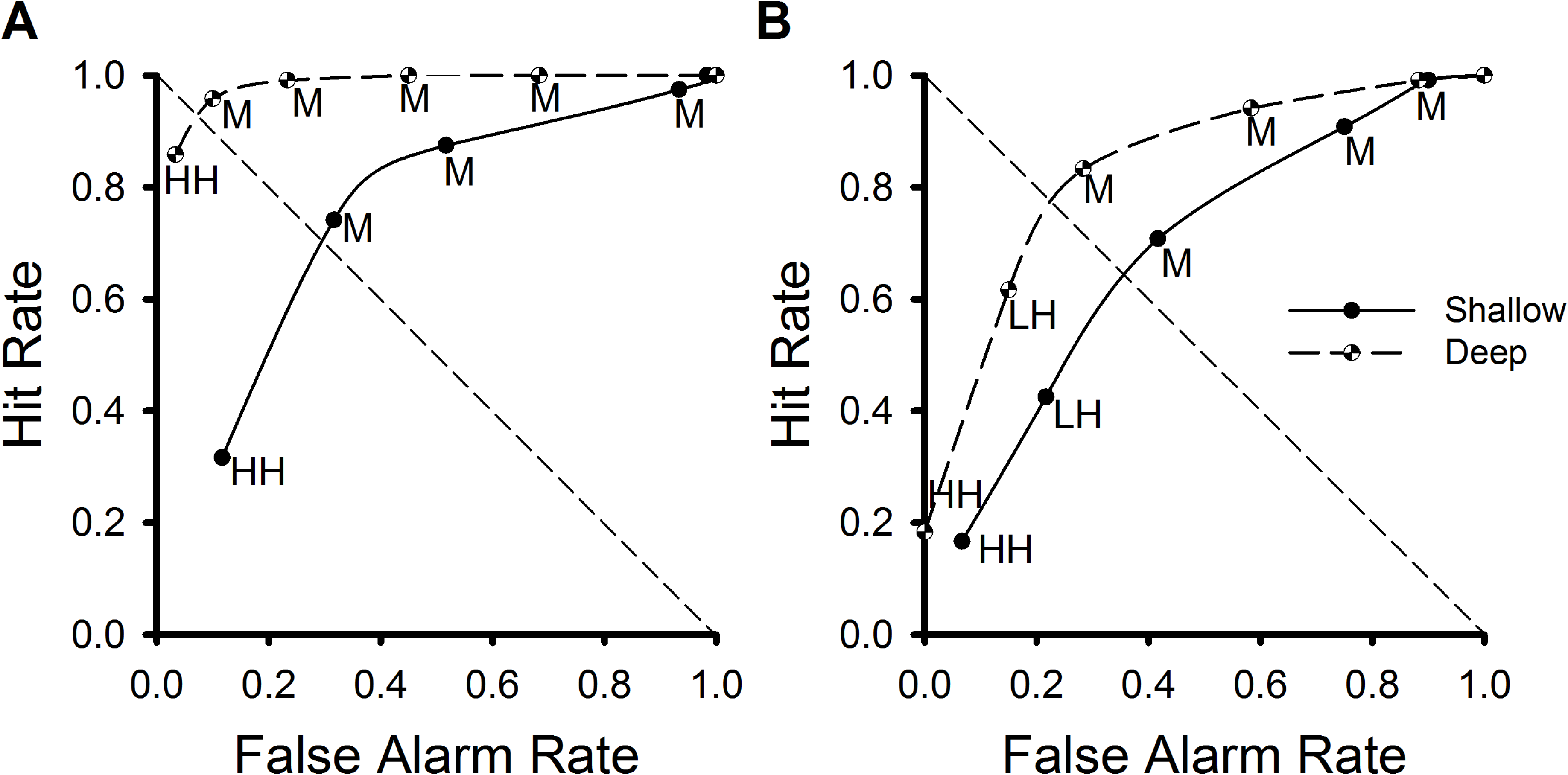
Receiver operating characteristic curves. ROC curves for a representative control (A) and Parkinson’s disease participant (B) in deep-semantic and shallow-non-semantic encoding conditions. The false alarm rate is cumulative. The responses given on the 6-point rating scale are grouped into the following conditions: high confidence hit (HH); low confidence hit (LH); miss (M).

Z scores were calculated for each individual using MATLAB (The Mathworks). Hits refer to combined HH and LH responses when a word is correctly remembered. False alarms are responses where the participant has incorrectly identified a new word as remembered.

### EEG data analysis

All EEG analysis and presented data are from the *encoding stage*. Offline analysis was performed in MATLAB using the open-source FieldTrip toolbox (Oostenveld, Fries, Maris, & Schoffelen, 2011) and in-house MATLAB functions. Raw EEG data were highpass (1 Hz) and lowpass filtered (40 Hz) with finite impulse response filters, re-referenced to the average reference, down-sampled to 500 Hz and epoched into 7000 ms segments around word presentation (3000 ms pre to 4000 ms post stimulus onset) for pre-processing. Independent component analysis allowed components related to ocular artefacts to be visually identified and removed before subsequent visual inspection and manual removal of remaining artefacts. If any channels had been removed during artefact rejection (mean of 0.6 channels removed, min: 0, max: 3), sensor data were interpolated via triangulation of nearest neighbour and then finally re-referenced to the average reference.

The EEG recording epochs extracted from individual encoding trials were grouped into HH, LH and M categories, depending on the participant’s subsequent response in the *recognition stage*. Epochs were further segmented from 750 ms pre-stimulus to 2000 ms post stimulus for the time-frequency analysis. The entire power spectrum was corrected for 1/f (Podvalny et al., 2015; Voytek et al., 2015) by fitting a linear function to the log-transformed data for every time point and then subtracting the linear fit. The 2.75 s epochs were then subjected to a Morlet wavelet transformation (width of 7 cycles) as implemented in Fieldtrip to extract time-frequency characteristics at frequencies 2 – 40 Hz in steps of 1 Hz. Average power values were calculated for each trial type (HH, LH and M) and baseline corrected (relative change, baseline −750 to −250 ms). This baseline duration is common to examine beta ERD in memory paradigms (e.g. (Hanslmayr et al., 2009; Meconi et al., 2016)) and the timing avoids filter smearing from post stimulus effects into the baseline period. The primary dependent measure was beta power decrease (i.e. ERD) for words that were subsequently successfully remembered, regardless of confidence level (i.e. during successful encoding of a memory resulting in a HH or LH trial in the recognition stage). The analysis of beta ERD between and within groups included an average of 101 (min 66/max 118) trials for controls and 98 trials (min 66/max 118) for PD participants in deep-semantic encoding, and 71 trials for both controls (min 33/max 98) and PD participants (min 25/max 103) in shallow-non-semantic encoding. The secondary dependent measure was the subsequent-memory effect (SME) in beta power which compared power between HH and M trials (Brewer et al., 1998; Hanslmayr et al., 2009; Otten et al., 2001).

### Statistical analysis

For memory strength (d’), participants who had values outside 3 standard deviations of the group mean were removed using the median absolute deviation method. The Shapiro-Wilk test ensured normality before using a mixed-effects repeated-measures 2X2 analysis of variance (ANOVA) with factors Group (Controls, PD) and Encoding (Deep, Shallow) as per our pre-registered protocol expecting a Group X Encoding interaction (MacDonald H, Jenkinson N, Hanslmayr S. Memory encoding and beta desynchronisation in Parkinson’s disease [Internet]. 2016 Available from: https://osf.io/vb64n/). Post-hoc and planned comparisons were performed using *t*-tests. A least absolute shrinkage and selection operator (LASSO) regression was performed for the PD group to determine the capacity of age and/or disease duration to predict memory strength following deep-semantic and shallow-non-semantic encoding, accounting for collinearity between age and disease duration. A mixed-effects repeated-measures 2X2 ANOVA with factors Group (Controls, PD) and Encoding (Deep, Shallow) tested for differences between groups in encoding accuracy and reaction time for the two encoding conditions.

In alignment with previous EEG studies, and as per our pre-registered protocol, post-stimulus beta power decreases are expected to be associated with successful memory formation in healthy (Hanslmayr et al., 2009; Hanslmayr et al., 2011) and patient populations (Meconi et al., 2016). Therefore lower beta from 12 – 20 Hz was the main frequency range of interest for all dependent measures (see https://osf.io/vb64n/). Only negative clusters in this frequency range were expected so comparisons of scalp-wide group averaged data were subjected to one-tailed cluster-based permutation testing (2000 iterations) using the Monte-Carlo ‘maxsum’ method (Meconi et al., 2016), averaged over 12 – 20 Hz and 0 – 1.5 s relative to encoding stimulus onset. The time window of 0 – 1.5 s post encoding stimulus was chosen based on findings from previous studies investing beta ERD using the same or similar memory paradigm (Hanslmayr et al., 2009; Meconi et al., 2016) and to avoid capturing any motor-related beta activity prior to the cue for a motor response (Pfurtscheller & Lopes da Silva, 1999) which appeared at the end of the encoding period (2 s after encoding stimulus). Data from all 128 electrodes are included in all EEG analyses. The only exception is for the additional correlational analyses in PD patients to further investigate the effect of encoding on their beta ERD at an individual level, when a subset of only left frontal electrodes was used based on a literature-driven prior hypothesis (Hanslmayr et al., 2009; Hanslmayr et al., 2011; Meeuwissen et al., 2011). This subset consisted of the front left quadrant taken from left sagittal to vertex (D23 – A1 on BioSemi cap), and vertex down to mid frontal (A1 – C17). A 2x2 mixed-effects repeated-measures ANOVA also tested for an Encoding (Shallow, Deep) X Group (Controls, PD) interaction of beta ERD and SME averaged for each participant over 0 – 1.5 s, 12 – 20 Hz and significant cluster electrodes. Linear regression tested for a relationship between each PD individual’s maximum beta desynchronization over left frontal electrodes during deep-semantic encoding and i) memory strength and ii) disease duration.

The criterion for all statistical significance was α = 0.05. Greenhouse-Geisser P values are reported for non-spherical data.

### Data availability

Anonymized data, not published in the article, will be shared on reasonable request from a qualified investigator.

## Acknowledgements

We thank Federica Meconi, Danying Wang and Sophie Watson for assistance with data collection.

